## Supplemental figures for "Clr4^SUV39H1^ ubiquitination and non-coding RNA mediate transcriptional silencing via heterochromatic phase transitions" (from another version of this preprint: version 3, 2023-08-04)

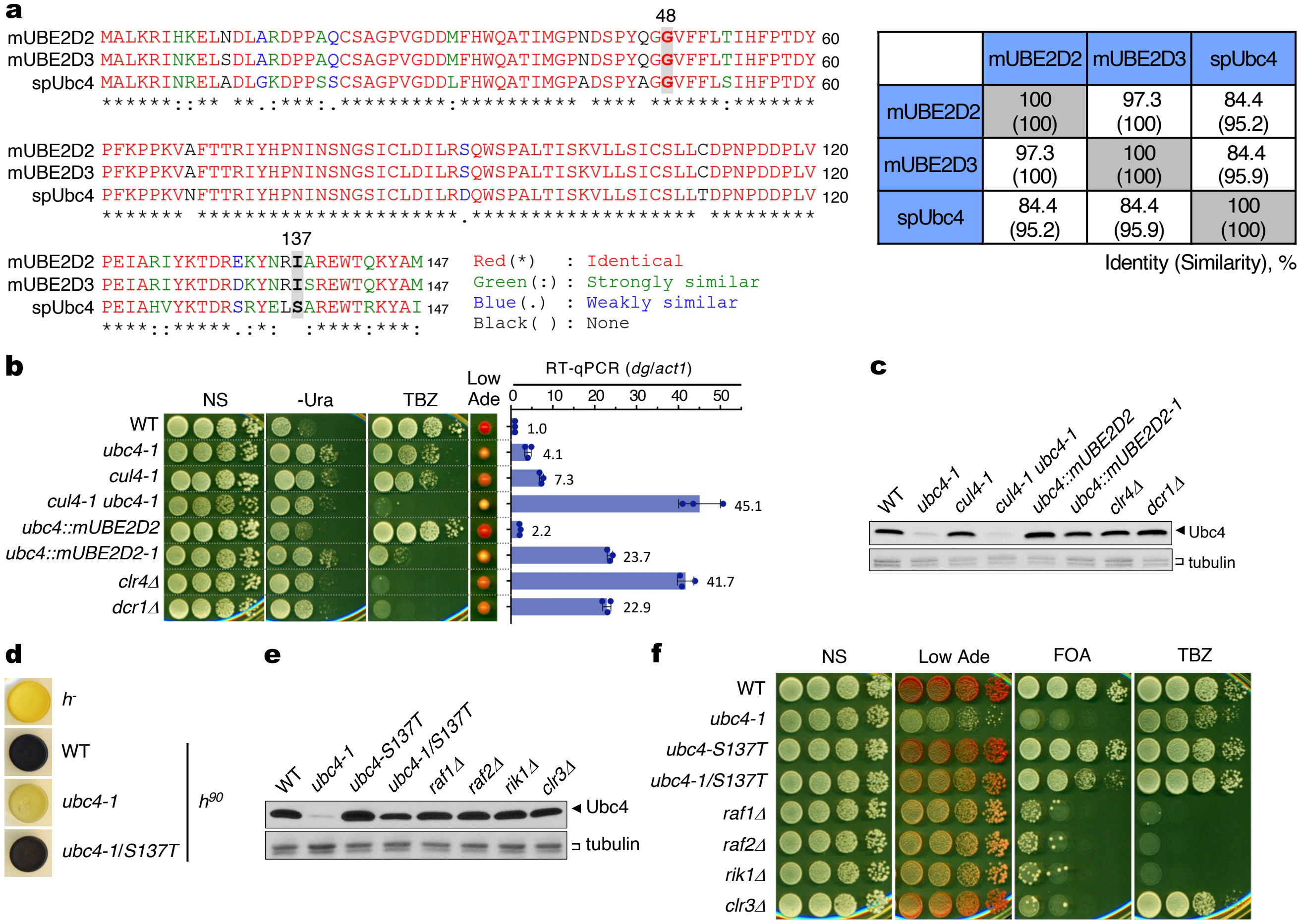


**Extended Data Fig. 1. Ubc4 and Cul4 regulate heterochromatin silencing.**
**a**, Left, Amino acid sequence comparison between Ubc4 with its mouse homologs using Clustal Omega. Glycine 48 of Ubc4 is conserved but Serine 137, which is important for Ubc4 stability (see below), is not conserved. Right, the summary of amino acid sequence comparison between *S. poombe* Ubc4 with mouse UBE2D2 and UBE2D3.
**b**, Assays for silencing of centromeric *ura4+* and *ade6+* reporter genes on no Uracil (-Ura) and low-adenine (Low Ade) media, and for chromosome segregation by viability on 15 µg/ml thiabendazole (TBZ) medium were performed in the indicated strains. Colonies turn to red when *ade6+* is silent. NS, non-selective medium. Right, RT-qPCR analysis for centromeric *dg* repeat (normalized to *act1*). Error bars, S.D. (n=3). The means are indicated on top of bars.
**c** and **e**, Ubc4 protein was greatly reduced in *ubc4-G48D* (*ubc4-1)* mutant. For WB analyses of Ubc4 protein in the indicated strains, antibody for mUBE2D3 was used to detect Ubc4. WB of tubulin was used as loading control.
**d**, Mating-type switching defect in *ubc4-1* mutant was suppressed by the secondary mutation, *S137T*. Non-switchable *h-* and WT and mutants in switchable strain (*h90*) grown on sporulation medium were stained with iodine vapors to test switch efficiency and spore formation. Dark color shows switch efficiency and pale color shows switch deficiency.
**f**, Assays for silencing of centromeric *ade6+* and *ura4+* on Low Ade and FOA media, a toxigenic substrate for Ura4, and for chromosome segregation by viability on 15µg/ml TBZ medium in the indicated strains.

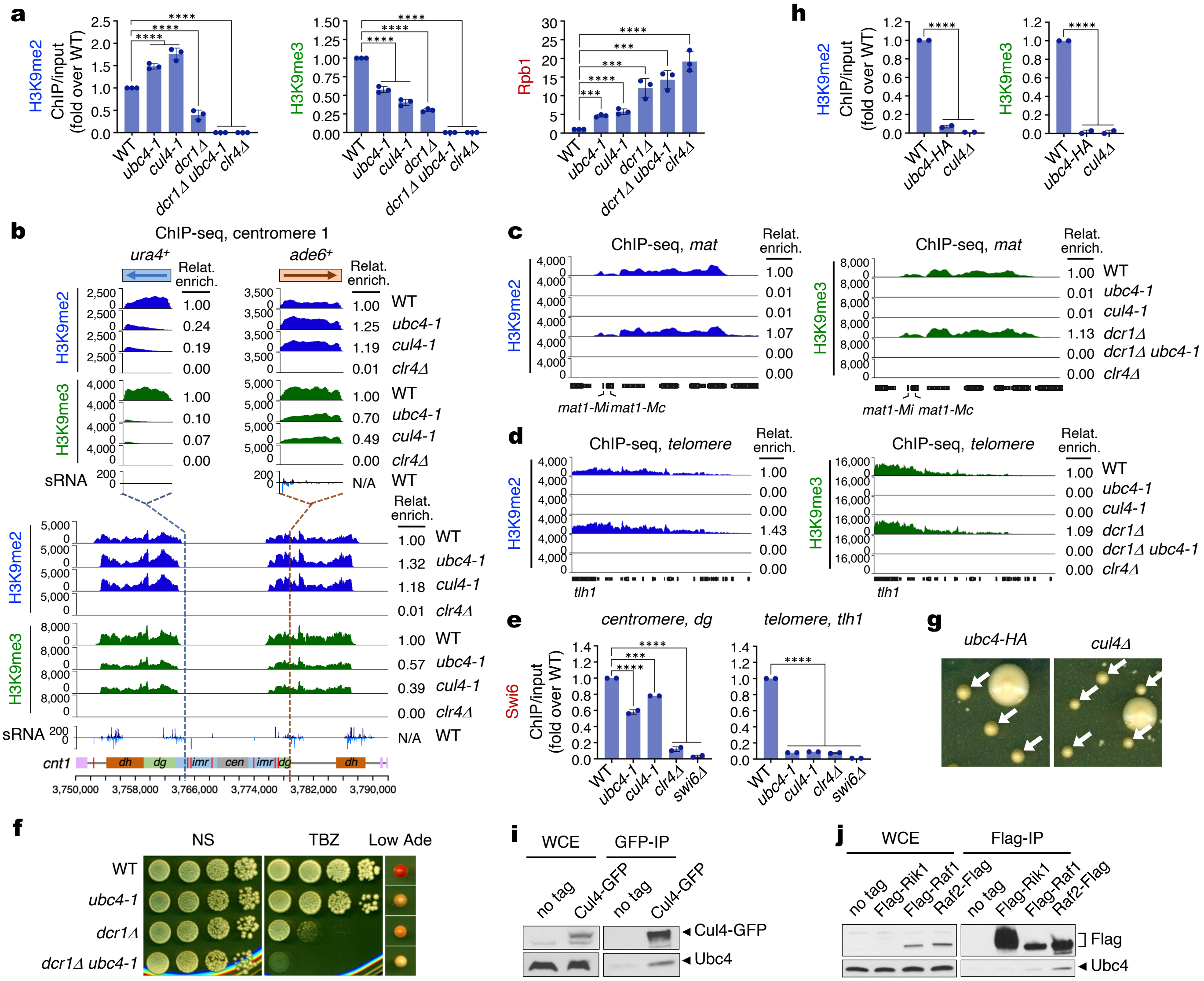


**Extended Data Fig. 2. Ubc4 and Cul4 regulate heterochromatin silencing by forming Ubc4-CLRC.**

**a**-**c**,ChIP-qPCR assays showing enrichment of H3K9me2 (**a**), H3K9me3 (**b**) and Rpb1 (**c**) at centromeric *dh* locus in the indicated strains. Error bars, S.D. (n = 3). *P* values are from Dunnett's multiple comparisons tests. $P < 0.0001$ for all mutants (**a** and **b**). $P = 0.0002$ for *ubc4-1*, $P < 0.0001$ for *cul4-1*, $P = 0.0008$ for *dcr1Δ*, $P = 0.0003$ for *dcr1Δ ubc4-1* and $P < 0.0001$ for *clr4Δ* (**c**).

**b**-**d**, ChIP-seq reads of H3K9me2 and H3K9me3 mapped to chromosome 1 centromere (**a**), mating type locus (**b**) and telomere (**c**) in the indicated strains. The change of H3K9me2 and H3K9me3 on *ura4+* and *ade6+* reporter genes and sRNA-seq reads of WT are inset (**a**). N/A, not available.

**e**, ChIP-qPCR assays showing enrichment of HP1 protein, Swi6 at centromeric *dg* locus (left) and telomere 1 left region (right) in the indicated strains. Error bars, S.D. (n = 2). *P* values are from Dunnett's multiple comparisons tests. $P < 0.0001$ for *ubc4-1*, $P = 0.0003$ for *cul4-1* and $P < 0.0001$ for *clr4Δ* and *swi6Δ* (left). $P < 0.0001$ for all mutants (right).

**f**, Assays for silencing of centromeric *ade6+* on Low Ade medium and for chromosome segregation by viability on 12.5 μg/ml TBZ medium in the indicated strains.

**g**, *ubc4* mutants tagged with triple HA at carboxyl-terminus (*ubc4-HA*) and *cul4* deletion mutants (*cul4Δ*) are viable but have a severe growth defect (arrows). Large colonies are WT controls.

**h**, ChIP-qPCR assays showing enrichment of H3K9me2 and H3K9me3 at centromeric *dg* repeat in the indicated strains. Error bars, S.D. (n = 2). *P* values are from Dunnett's multiple comparisons tests. $P < 0.0001$ for all mutants (left and right).

**i** and **j**, Co-immunoprecipitation (Co-IP) of Cul4-GFP (**H**) and Flag-Rik1, Flag-Raf1 and Raf2-Flag (**I**) with Ubc4.

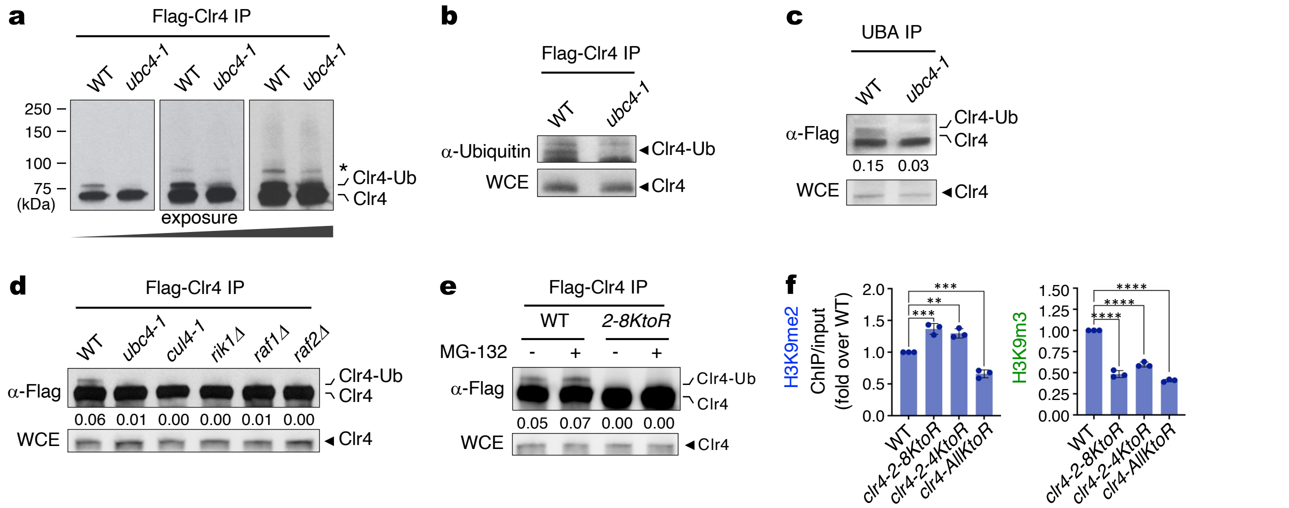


**Extended Data Fig. 3. Mono-ubiquitination of Clr4.**

**a**, Extended exposure of WB analysis for Flag-Clr4 IP from WT and *ubc4-1* mutant cells (Fig. 1B). Asterisk indicates possible di- ubiquitination which is also largely dependent on Ubc4.

**b**, Flag-Clr4 IP and WB analysis using ubiquitin antibody.

**c**, WB analyses Flag-Clr4 enrichment by ubiquitination affinity beads (UBA IP).

**d**, WB analyses of immunoprecipitated Flag-Clr4 (Flag-Clr4 IP) in the indicated strains.

**e**, Proteasome-independent Clr4 ubiquitination. WB analyses of Flag-Clr4 IP with or without treatment of proteasome inhibitor MG-132 in the indicated strains.

**f**, ChIP-qPCR assays showing enrichment of H3K9me2 and H3K9me3 at centromeric *dh* repeat in the indicated strains. Error bars, S.D. (n = 2). *P* values are from Dunnett's multiple comparisons tests. $P < 0.0001$ for all mutants (left and right).

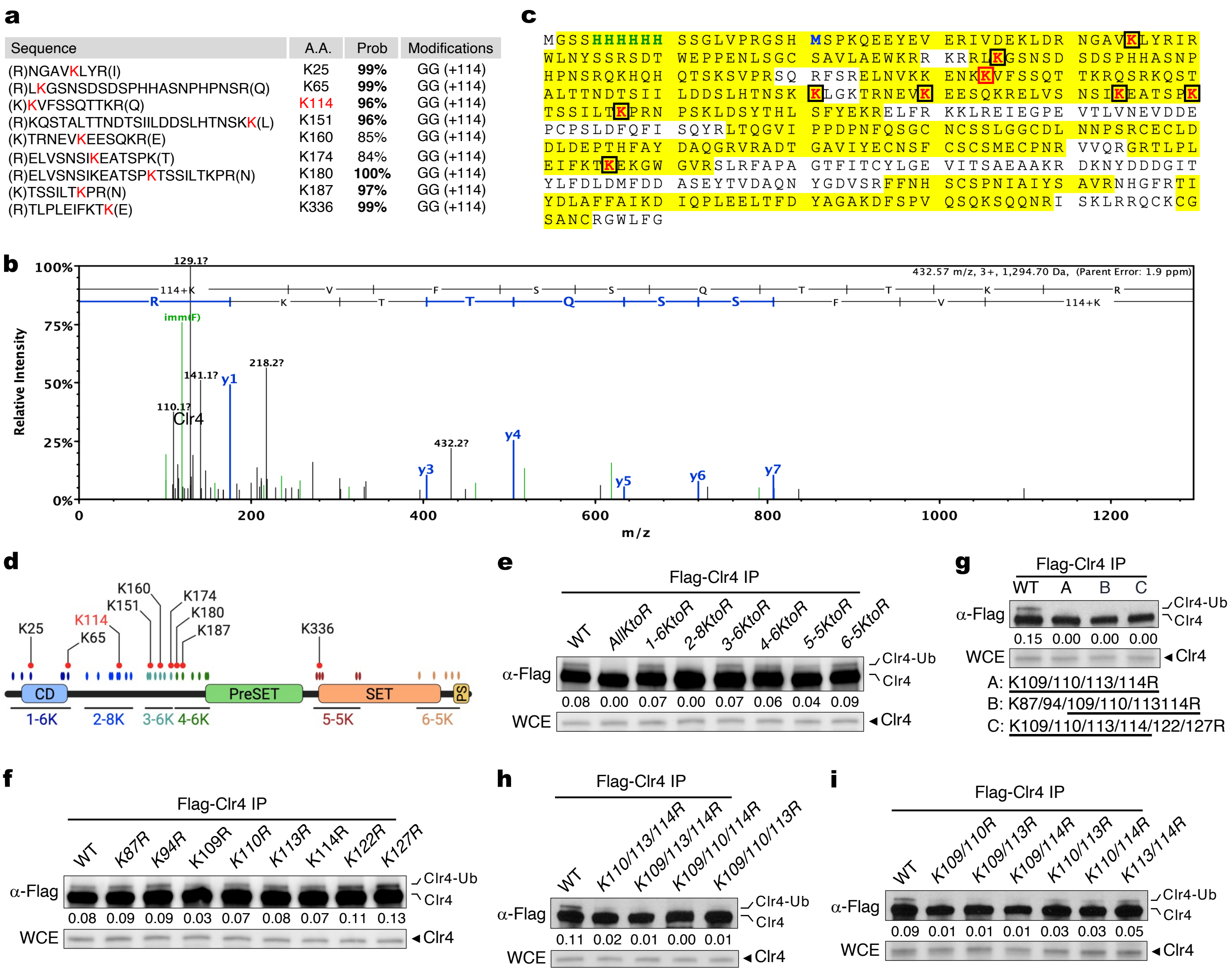


**Extended Data Fig. 4. Characterization of ubiquitination sites of Clr4 by Mass Spec analysis.**

**a**, 6xHis-Clr4 purified from *E. coli* cells were *in vitro* ubiquitinated and analyzed by Mass Spectrometry. Di-glycine (GG) remnants after trypsin digestion indicate possible ubiquitination sites. Prob., probability. K114 is highlighted.

**b**, MS/MS spectrum of the peptide containing ubiquitinated K114. The observed y ions are shown. Predicted amino acids sequences are across the top.

**c**, Amino acid sequence of 6xHis-Clr4 protein. Sequence coverage is highlighted in yellow. 6xHis (green) and translation start site (blue) are shown. Possible ubiquitination sites (red) are highlighted with black boxes and K114 is highlighted with red box.

**d**, Schematic diagram of domain structure of Clr4 and ubiquitination sites characterized by Mass Spectrometry analysis. K114 is highlighted with red.

**e**-**i**, WB analyses of Flag-Clr4 IP from indicated strains.

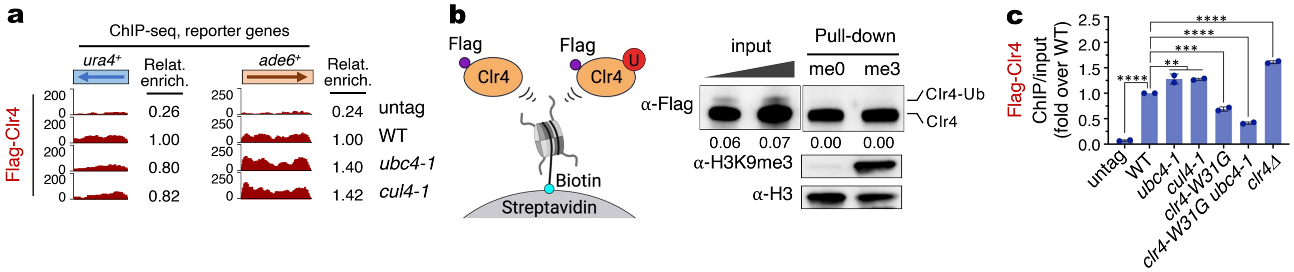


**Extended Data Fig. 5. Clr4 mono-ubiquitination induces its dissociation from chromatin and from centromeric ncRNA.**

**a**, ChIP-seq reads of Flag-Clr4 mapped to *ura4+* and *ade6+* reporter genes in indicated strains.

**b**, Binding assay of Flag-Clr4 protein purified from *S. pombe* cells with nucleosomes with unmodified histone H3 (me0) or H3K9me3 histone (me3) anchored to streptavidin beads. Input and Flag-Clr4 proteins from pull-down were analyzed by WB using antibodies as shown. Schematic diagram of binding assay (left).

**c**, ChIP-qPCR assays showing enrichment of Flag-Clr4 at centromeric *dh* repeat in the indicated strains. Error bars, S.D. (n = 2). *P* values are from Dunnett's multiple comparisons tests. $P < 0.0001$ for untag, $P = 0.0017$ for *ubc4-1*, $P = 0.0018$ for *cul4-1*, $P = 0.0009$ for *clr4-W31G* and $P < 0.0001$ for *clr4-W31G ubc4-1* and *clr4Δ.*

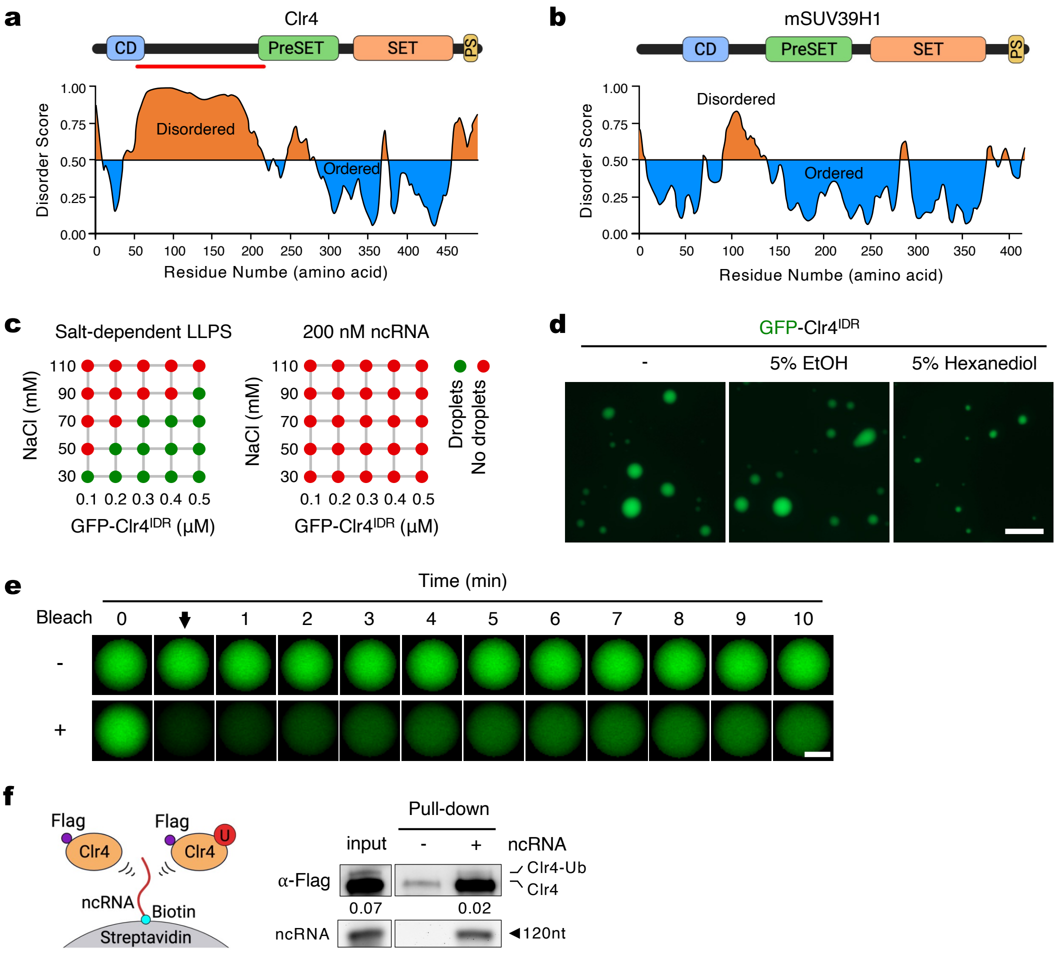


**Extended Data Fig. 6. Phase separation of Clr4.**

**a**, Schematic diagram of domain structure of Clr4 protein. Disorder prediction shows that the hinge region between the chromodomain (CD) and PreSET domain of Clr4 is highly disordered. Red bar shows the location of amino acids which is used for GFP-Clr4$^{IDR}$ fusion (51-220 amino acids).

**b**, Schematic diagram of domain structure and disorder prediction (PONDR) of mSUV39H1 protein.

**c**, Summary of LLPS behaviors of Clr4$^{IDR}$ with different concentrations of salt (NaCl, left). Summary of LLPS behaviors of Clr4$^{IDR}$ with different concentrations of salt in the presence of 200 nM centromeric ncRNA (right).

**d**, Representative images of phase-separated liquid droplets of GFP-Clr4$^{IDR}$ after treatment with 5% Ethanol (EtOH) or 5% Hexanediol. Scale bar, 5 µm.

**e**, FRAP of GFP-Clr4$^{IDR}$. Representative phase-separated liquid droplests were monitored as shown. Arrow shows the time of bleaching (~40s). Scale bar, 2 µm.

**f**, Flag-Clr4 protein purified from *S. pombe* cells were incubated with centromeric ncRNA anchored to streptavidin beads. Input and bound Flag-Clr4 proteins from pull-down were analyzed by WB. ncRNA from input and pull-down was stained with SYBR Gold. Schematic diagram (left).

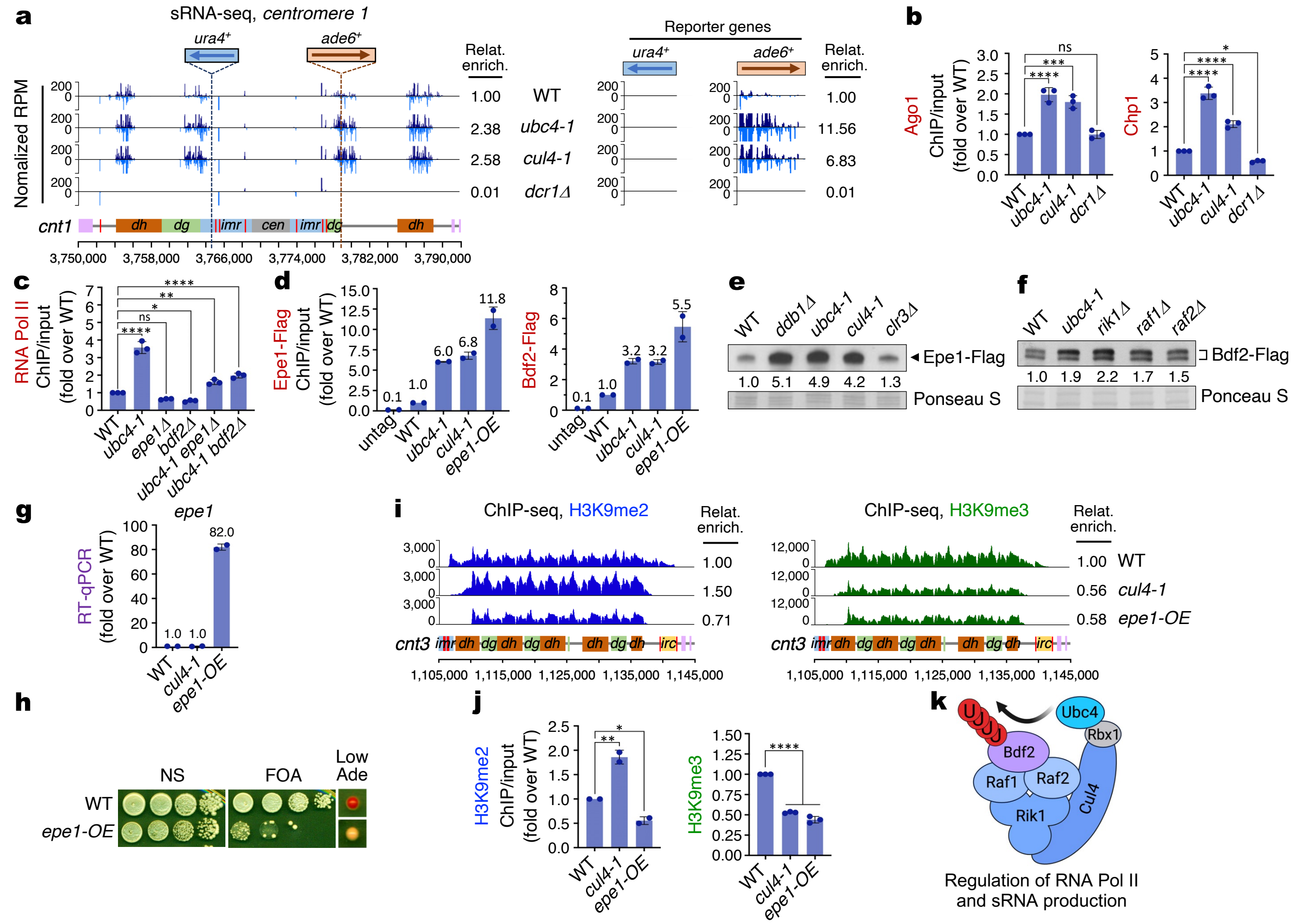


**Extended Data Fig. 7. Ubiquitination by Ubc4 and Cul4 regulates centromeric sRNA transcription through Epe1 and Bdf2$^{BRD4}$.**

**a**, sRNA-seq reads mapped to chromosome 1 centromere. The *ura4+* reporter gene is inserted in *imr* with no sRNA production whereas *ade6+* is located in *otr* where sRNA are produced. Increased sRNA production was observed at the *ade6+* reporter gene, but not at the *ura4+* reporter gene, in *ubc4-1* and *cul4-1* mutants.

**b**, ChIP-qPCR assays for enrichment of Ago1 and Chp1 at centromeric *dh* repeat in the indicated strains. Error bars, S.D. (n = 3). *P* values are from Dunnett's multiple comparisons tests. $P < 0.0001$ for *ubc4-1* and $P = 0.0002$ for *cul4-1* (left). $P < 0.0001$ for *ubc4-1* and *cul4-1* and $P = 0.0206$ for *dcr1Δ.* (right).

**c**, ChIP-qPCR assays for enrichment of RNA Pol II (Rpb1) at centromeric *dg* repeat in the indicated strains. Error bars, S.D. (n = 3). $P < 0.0001$ for *ubc4-1*, $P = 0.0172$ for *bdf2Δ*, $P = 0.0028$ for *ubc4-1 epe1Δ* and $P < 0.0001$ for *ubc4-1 bdf2Δ*.

**d**, ChIP-qPCR assays for enrichment of Epe1-Flag and Bdf2-Flag at centromeric *dh* repeat in the indicated strains. Error bars, S.D. (n = 2).

**e** and **f**, WB analyses for Epe1-Flag and Bdf2-Flag in the indicated strains. Ponceau S staining is used as control.

**g**, RT-qPCR analysis for *epe1* transcripts in the indicated strains. Error bars, S.D. (n = 3).

**h**, Assays for silencing of centromeric *ura4+* and *ade6+* reporter genes on FOA and Low Ade media in the indicated strains.

**i**, ChIP-seq reads of H3K9me2 (blue) and H3K9me3 (green) mapped to centromere 3 (*cnt3*) right arm in the indicated strains.

**j**, ChIP-qPCR assays for enrichment of H3K9me2 and H3K9me3 at centromeric *dh* repeat in the indicated strains. Error bars, S.D. (n = 2). $P = 0.0045$ for *cul4-1* and $P = 0.0287$ for *epe1-OE* (left). $P < 0.0001$ for *cul4-1* and *epe1-OE* (right).

**k**, Schematic diagram showing the presumptive subunit arrangement in the complex of Ubc4-CLRC and its substrate Bdf2. Poly-ubiquitination activity of Ubc4-CLRC toward Bdf2 promotes degradation of Bdf2, reduced RNA Pol II recruitment to centromeric heterochromatin and subsequently reduced centromeric sRNA production. Red circle (U), ubiquitin.

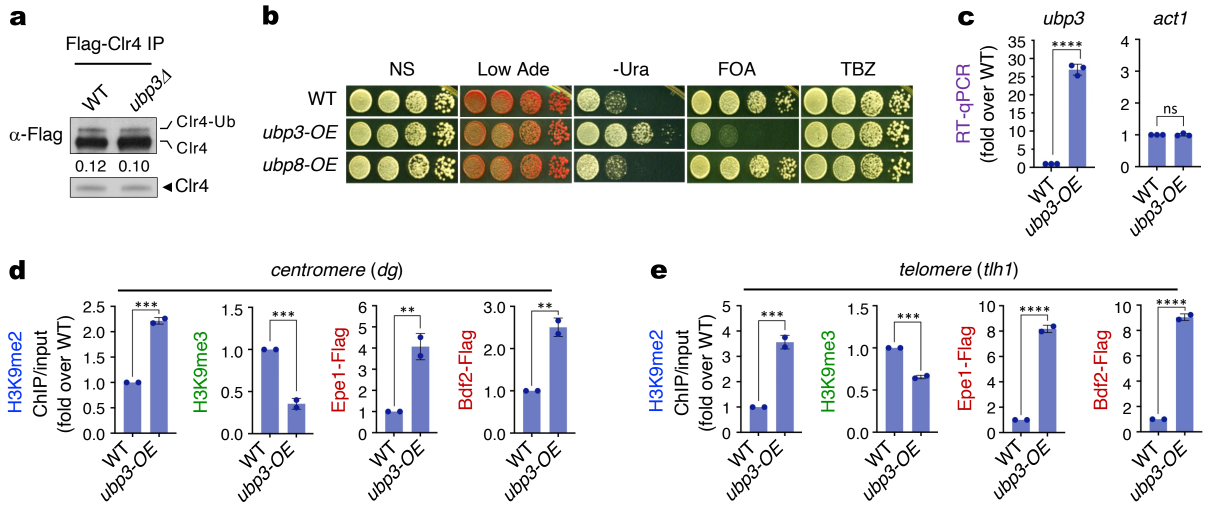


**Extended Data Fig. 8. Ubp3 counteracts Ubc4-CLRC activities.**

**a**, WB analyses of Flag-Clr4 IP reveals that *ubp3* deletion does not cause significant change of Clr4 ubiquitination.

**b**, Assays for silencing of centromeric $ade6^{+}$ and $ura4^{+}$ reporter genes on Low Ade, -Ura and FOA media and for chromosome segregation by viability on 15 µg/ml TBZ medium in *ubp3* or *ubp8* overexpression (-OE) cells.

**c**, RT-qPCR analysis for *ubp3*, centromeric *dg* and *act1* transcripts in the indicated strains. Error bars, S.D. (n = 3). $P$ values are from Dunnett's multiple comparisons tests. $P < 0.0001$ for *ubp3-OE* (left).

**d**, ChIP-qPCR assays for enrichment of H3K9me2, H3K9me3, Epe1-Flag and Bdf2-Flag at centromeric *dg* locus. Error bars, S.D. (n = 2). $P$ values are from Dunnett's multiple comparisons tests. $P = 0.0001$ for *ubp3-OE* (H3K9me2), $P = 0.0007$ for *ubp3-OE* (H3K9me2), $P = 0.0056$ for *ubp3-OE* (Epe1-Flag) and $P = 0.0021$ for *ubp3-OE* (Bdf2-Flag)

**e**, ChIP-qPCR assays for enrichment of H3K9me2, H3K9me3, Epe1-Flag and Bdf2-Flag at telomeric *tlh1* locus. Error bars, S.D. (n = 2). $P$ values are from Dunnett's multiple comparisons tests. $P = 0.0008$ for *ubp3-OE* (H3K9me2), $P = 0.0002$ for *ubp3-OE* (H3K9me2) and $P < 0.0001$ for *ubp3-OE* (Epe1-Flag) and *ubp3-OE* (Bdf2-Flag).

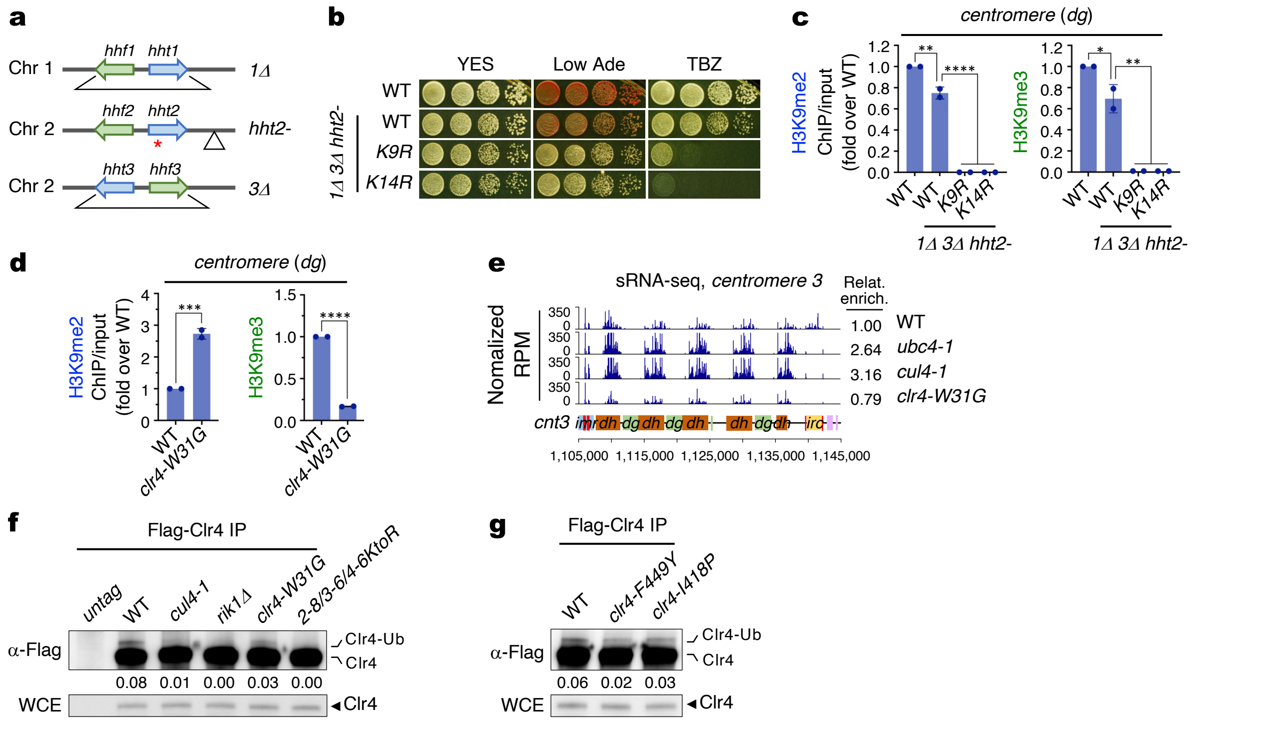


**Extended Data Fig. 9. Mutation of histone H3 lysines 9 and 14 abolishes both H3K9me2 and H3K9me3, while *clr4* mutants with H3K9me2-to-H3K9me3 transition defect show reduced mono-ubiquitination.**

**a**, Schematic diagram of histone H3 and H4 genes. After the *hhf1-hht1* and *hht3-hhf3* gene pairs were deleted (open trapezoid, *1Δ 3Δ*), the *hht2* gene was mutagenized by inserting a marker gene (triangle) and simultaneous homologous recombination (*hht2*-). Asterisk indicates mutation site in *hht2*.

**b**, Assays for silencing of centromeric *ade6*$^{+}$ on Low Ade medium and for chromosome segregation by viability on 15 µg/ml TBZ medium in the indicated strains. Both H3K14R and H3K9R mutants have a strong silencing defect with white colonies on Low Ade medium, and high sensitivity to TBZ.

**c**, ChIP-qPCR assays for enrichment of H3K9me2 and H3K9me3 at centromeric *dg* repeats in the indicated strains. Error bars, S.D. (n = 2). Both H3K14R and H3K9R mutants completely lost H3K9me2 and H3K9me3 at the centromere. *P* values are from Dunnett's multiple comparisons tests. $P = 0.0021$ for *1Δ 3Δ hht2*-WT and $P < 0.0001$ for *1Δ 3Δ hht2-K9R and 1Δ 3Δ hht2-K14R* (left). $P = 0.0238$ for *1Δ 3Δ hht2*-WT and $P = 0.0012$ for *1Δ 3Δ hht2-K9R and 1Δ 3Δ hht2-K14R* (right).

**d**, ChIP-qPCR assays showing enrichment of H3K9me2 (left), H3K9me3 (right) at centromeric *dg* repeat in the indicated strains. Error bars, S.D. (n = 2). *P* values are from Dunnett's multiple comparisons tests. $P = 0.0006$ for *clr4-W31G* (left) and $P < 0.0001$ for *clr4-W31G* (right).

**e**, sRNA-seq reads mapped to centromere 3 right arm in indicated strains.

**f** and **g**, WB analyses of Flag-Clr4 IP from indicated strains.

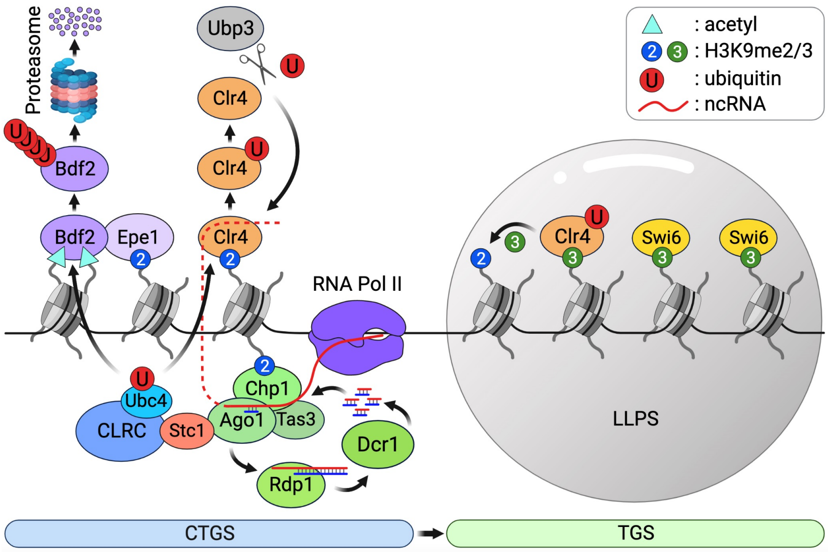


**Extended Data Fig. 10. Model for heterochromatin phase transition.**
The histone H3 lysine-9 (H3K9) methyltransferase Clr4$^{SUV39H1}$ is recruited to chromatin by interactions with methylated H3K9 and with non-coding RNA, which is processed by RNAi (dotted red line). Ubc4-CLRC is recruited by RNAi, and mono-ubiquitinates Clr4$^{SUV39H1}$, releasing it from chromatin and from non-coding RNA. Release promotes heterochromatic phase separation along with Swi6, and the transition from H3K9me2 (blue) to H3K9me3 (green). Ubc4-CLRC also poly-ubiquitinates Bdf2$^{BRD4}$, a transcriptional co-activator required for Pol II transcription and small RNA production. In this way, Ubc4-CLRC mediates the transition from co-transcriptional gene silencing (CTGS), mediated by RNAi, to transcriptional gene silencing (TGS) mediated by phase separation of H3K9me3 and Swi6$^{HP1}$.
